# Spatial distribution and hotspots of mammals in Canada

**DOI:** 10.1101/2020.01.29.925461

**Authors:** Victor Cameron, Anna L. Hargreaves

**Author notes:** **Correspondent**: Anna L. Hargreaves –.

## Abstract

High-latitude countries often contain the polar range edge of species that are common farther south. The more peripherally a species occurs in a country, the smaller its national range will be and the more its national range will consist of range-edge populations, which are often predicted to be relatively small, isolated, and unproductive. Together, this may focus national conservation efforts toward peripheral species whose global conservation value is controversial. However, if range-edge taxa occur where overall diversity is also high, there would be fewer trade-offs in protecting them. Using 153 of the 158 terrestrial mammal species in Canada, we tested how species’ distributions relate to their national conservation status and total mammal richness. Half of ‘Canadian’ mammals had <20% of their global range in Canada. Range area in Canada was strongly associated with national threat status; mammals considered ‘at-risk’ in Canada had 42% smaller Canadian ranges than mammals considered secure. However, after accounting for range area, being more peripheral (smaller proportion of global range in Canada) did not increase the likelihood that a taxon was considered at-risk. We overlaid the 153 maps to calculate mammal diversity across Canada, divided into 100×100 km grid cells. We found that hotspots of at-risk mammals (cells with >4 at-risk taxa) and hotspots of range-edge mammals (cells with >12 taxa with ≤20% of their range in Canada) were about twice as species rich as non-hotspot cells, containing up to 44% of Canadian mammal diversity per grid-cell. Our results suggest that protecting areas with the most at-risk or range-edge mammals would simultaneously protect habitat for many species currently deemed secure.

## Introduction

Large polar countries often contain the high-latitude range edge of many species that are more widely distributed beyond their borders (Cheffings et al. 2005; Gibson et al. 2009; Rassi et al. 2010; ArtDatabanken 2015). For a given species, the more peripherally it occurs in a country, the smaller its national range area will be. Thus range-edge taxa that occupy small areas in a country are more likely to be deemed nationally at-risk (Lesica & Allendorf 1995), especially if range-edge populations themselves are smaller, more isolated, or less productive than more central populations (Brown et al. 1996; Sagarin & Gaines 2002; Yakimowski & Eckert 2007; although this pattern is far from universal; Samis & Eckert 2009; Pironon et al. 2017; Hargreaves & Eckert 2019). Conservation in large polar countries may therefore be focused toward range-edge species even for taxa that are globally secure (Hunter & Hutchinson 1994).

The value and practicality of conserving range-edge populations are contentious. Ethically, some argue that countries have the greatest obligations to taxa with the largest percentage of their range in their borders, and that taxa are not truly at-risk if they are globally secure (Hunter & Hutchinson 1994). Practically, edge populations may be difficult to conserve if they are inherently unstable due to small numbers, low genetic diversity, or poor habitat (Hunter & Hutchinson 1994; although again these patterns do not hold for many taxa; Eckert et al. 2008). On the other hand, range-edge populations can be important for diversification if they occupy unique habitats (Van Rossum et al. 2003; Mimura & Aitken 2010), and natural range expansion following gradual local adaptation (Hargreaves & Eckert 2019). Polar edge populations are also geographically poised to initiate range shifts in response to climate warming (Gibson et al. 2009). The geographic ‘head start’ edge populations offer is especially important when species’ dispersal ability is low compared to the rate of climate change, as is the case for many terrestrial mammals (Thomas et al. 2004; Schloss et al. 2012).

Evaluating the relative conservation merits of range-edge taxa would be less important if protecting edge populations involved few conservation trade-offs. One important conservation strategy is establishing protected areas (Myers et al. 2000; Le Saout et al. 2013). Protected areas will be most effective if multiple levels of biodiversity co-occur (Prendergast et al. 1993; Myers et al. 2000; Ricketts et al. 2005), e.g. if at-risk or range-edge species occur in areas with high species richness. Previous studies have not found significant co-occurrence of rare or threatened taxa and overall species richness, either at a global scale for vertebrates (Orme et al. 2005; Ceballos & Ehrlich 2006; Jenkins et al. 2013), or within the UK for invertebrates, plants and birds (Prendergast et al. 1993). However, co-occurrence may be more likely in large high-latitude countries if range-edge taxa are often considered nationally at-risk, since edge taxa and biodiversity will be concentrated toward the equatorward border of such countries (Buckley et al. 2010).

We explore the co-occurrence of at-risk and range-edge taxa with overall richness using terrestrial mammals in the world’s second largest country: Canada. Canadian mammals provide an excellent case study as distribution maps and national threat ranks are available for each taxon (usually species but sometimes subspecies or populations), and Canada’s conservation assessment body–the Committee on the Status of Wildlife in Canada (COSEWIC)–uses IUCN criteria that emphasize local abundance and range area. Protection of edge populations in Canada is also of global conservation significance, as Canada contains the highest latitude refuges for species in the Americas. Many species designated ‘at-risk’ in Canada are range-edge populations of species largely distributed outside Canada (Gibson et al. 2009; Klemet-N’Guessan et al. 2019), but previous estimates have not compared at-risk to secure taxa or quantified range areas in Canada. It is therefore unclear whether so many at-risk taxa are peripheral simply because most taxa in Canada are peripheral, because edge populations have smaller range areas in Canada, or because edge populations are inherently more at risk.

Using IUCN range maps available for 153 of the 158 extant mammal species that occur in Canada, and categorizing each taxon as nationally at-risk or secure based on its COSEWIC status, we asked two questions. *Question 1*) How do range area in Canada and the proportion of a taxon’s global range in Canada (lower means more peripheral) relate to national conservation status? We predicted that: a) at-risk taxa will have smaller Canadian ranges given the link between range area and population size (Lawton 1993) and importance of both for conservation; b) at-risk taxa will have smaller percentages of their range in Canada given previous findings that many at-risk taxa are edge populations (Gibson et al. 2009; Klemet-N’Guessan et al. 2019); and c) that smaller range percentages would increase the probability that a taxon is at risk even after controlling for range area, if edge populations routinely suffer from factors that would increase their conservation risk (Coristine & Kerr 2011). We then divided Canada into 100×100 km grid-cells and asked *Question 2*) Do cells with the high richness of at-risk species or range-edge species (hotspots) occur in regions of high overall mammal richness? If range-edge populations are over-represented among at-risk species, as predicted in Question 1, and since edge populations and species richness should both cluster toward Canada’s low-latitude border, we predict that at-risk and range-edge hotspots will have higher total mammal richness than non-hotspot cells.

## Methods

We recorded the Canadian conservation status for each mammal taxon from COSEWIC assessments; we used COSEWIC’s recommendation rather than official status under the Species at Risk Act (SARA) as COSEWIC addresses only conservation risk whereas SARA also considers the economic impact of a given listing. Taxa per threat category: 10 Special concern, 5 Threatened, 7 Endangered, 8 assessed as Not at Risk. The remaining 123 mammal taxa had not been assessed. As COSEWIC assesses species in order of perceived risk, we assume that those that have not been assessed in the 41 years of COSEWIC assessments are relatively secure. Given the low sample sizes within at-risk categories, we binned taxa as either ‘at-risk’ (Special Concern, Threatened, or Endangered) or ‘secure’ (assessed as Not at Risk or not assessed).

We obtained distribution maps in the form of spatial polygons from the International Union for Conservation of Nature (IUCN 2018). We first selected the 158 terrestrial mammal species whose distribution polygon overlapped a polygon map of Canada (Natural Earth 2018). IUCN maps are by species, but COSEWIC sometimes assesses subspecies or populations. We retained subspecies and populations if the distribution map provided in their COSEWIC assessment was similar to their IUCN distribution polygon (2 subspecies and 1 population), but discarded 5 at-risk taxa whose IUCN polygons did not match the geographic scale of their COSEWIC assessments. This yielded 153 taxa: 150 species, 2 subspecies, and 1 population (153 unique species in total; Fig. 1). Each map was cropped to land only by overlapping it with a world boundary map (Natural Earth 2018) using the “sf” package (version 0.7-3, Pebesma 2018) in the statistical platform R (version 3.5.1, R Core Team 2018). As we were interested in range area, polygons were projected into Albers equal area projection (Gibson et al. 2009).

**Fig. 1:**
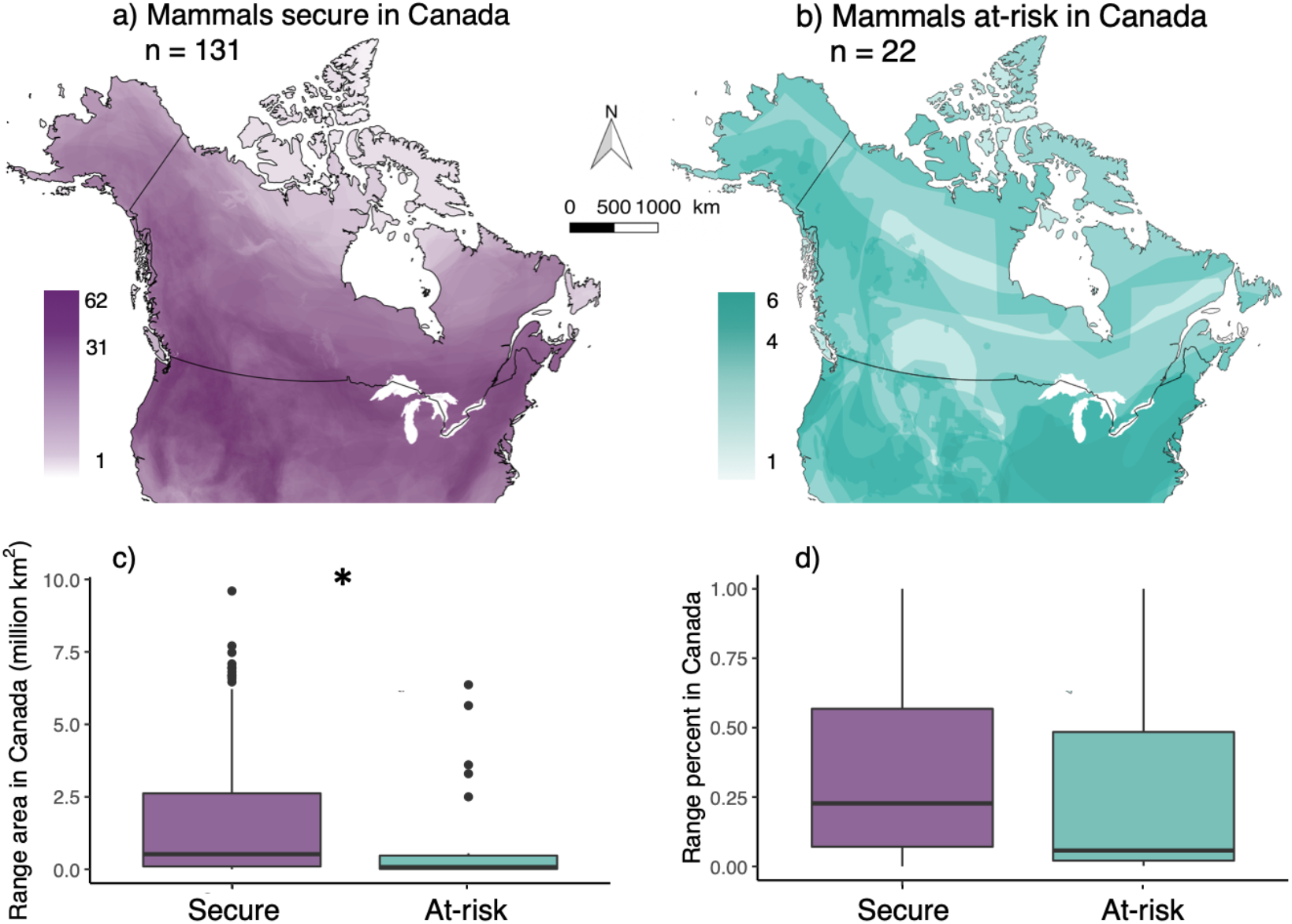
Geographic distributions of terrestrial mammals deemed secure (purple) and at risk (teal) in Canada. (a & b) Each polygon shows one taxon’s distribution, areas of darker colour contain more taxa (colour scale bars indicate species richness). Although only taxa that occur in Canada are mapped, areas of high richness occur south of the Canadian border. At-risk and secure mammals differ in Canadian range size (c; * indicates statistical difference from model considering ln-transformed data), but not the percentage of their global range that occurs in Canada (d). c and d show raw data, centre lines and box boundaries represent the median, 25^th^ and 75^th^ percentiles respectively, whiskers extend to 1.5x the distance between the 25^th^ and 75^th^ percentiles, dots are outliers.

### Q1) How do range area in Canada and range percentage in Canada relate to national conservation status?

We determined each taxon’s range area and range percentage in Canada using the “sf” package. We determined range area in Canada (km^2^) by overlapping each species’ map with a boundary map of Canada (Natural Earth 2018), then measuring the area of the overlap. Canadian range area varied from 2 km^2^ (*Lontra canadensis*) to >9 million km^2^ (*Nycticeius humeralis*), with mean = 1 678 850 km^2^ and median = 460 130 km^2^. Due to the large spread of range areas, we ln-transformed Canadian range area for analyses (see below). We quantified each taxon’s global range area by measuring the area of overlap between its range map and a global land map. Global range area varied from 2075 km^2^ (*Marmota vancouverensis*) to >22 million km^2^ (*Puma concolor*), with mean = 4 449 710 km^2^ and median = 3 263 110 km^2^. We calculated the percentage of each taxon’s global range that occurs in Canada (i.e. the inverse of peripherality in Canada) as Canadian range area/Global range area x 100; this varied from <0.001% (*Nycticeius humeralis*) to 100% (*Dicrostonyx nunatakensis* & *Marmota vancouverensis*).

We tested the relationships between range area in Canada, range percentage in Canada, and conservation status in Canada using three models. We tested whether range area (ln(range area), gaussian response; *model a*) or percentage (proportion response; *model b*) in Canada differed between taxa deemed at-risk or secure (conservation status = binomial predictor), using generalized linear models (GLMs) with a gaussian and binomial error structure, respectively (*glm* command, base R; ln(range area) or percentage ~ conservation status). Next, using a binomial GLM we tested whether the probability of a taxa being at-risk (conservation status = binomial response) varied with range percentage in Canada (proportion predictor) even after accounting for range size (continuous predictor; *model c*): conservation status ~ ln(range area) + range percentage (Venables & Ripley 2002). We assessed the significance of predictors using likelihood ratio Chisquare tests (*Anova* type=III command, *car* package).

### Q2) Do hotspots of at-risk or range-edge taxa coincide with high overall diversity?

To quantify the spatial distribution of mammal diversity, we overlaid a grid of 100×100 km cells on the Canadian equal area map. This produced a map of Canada divided into 1285 cells (not all equal in area as the Canadian border bisected some cells; Supporting Information). We overlaid the range map polygons of all terrestrial mammals on the grid map, then counted the total mammal species, at-risk species, and range-edge species in each cell. We defined ‘range-edge species’ as taxa with ≤20% of their global range in Canada (as per Klemet-N’Guessan et al. 2019); using a threshold of 10% reduced the number of range-edge taxa from 76 to 53 but did not alter conclusions to Questions 1 and 2 (Supporting Information). Following Prendergast et al. (1993) and Reid (1998), we identified cells with the highest richness (‘hotspots’) of at-risk and of range-edge taxa up to a maximum of 5% of the total grid cells. We use ‘hotspot’ *sensu* Prendergast et al. (1993) to refer to areas with high biodiversity, rather than *sensu* Myers (1988) as areas of high diversity *and* high extrinsic threat. This yielded 47 hotspot cells for at-risk taxa (5 to 6 at-risk taxa/cell; 3.6% of total grid cells; Fig. 2a), and 47 hotspot cells for range-edge taxa (17 to 29 range-edge taxa/cell; Fig. 2b). Cells containing 4 at-risk or 16 range-edge taxa were not counted as the number of hotspot cells would have exceeded 5% of total cells. This method of identifying cells of high richness creates richness indices that are reasonably independent of total species richness (Reid 1998).

**Fig. 2:**
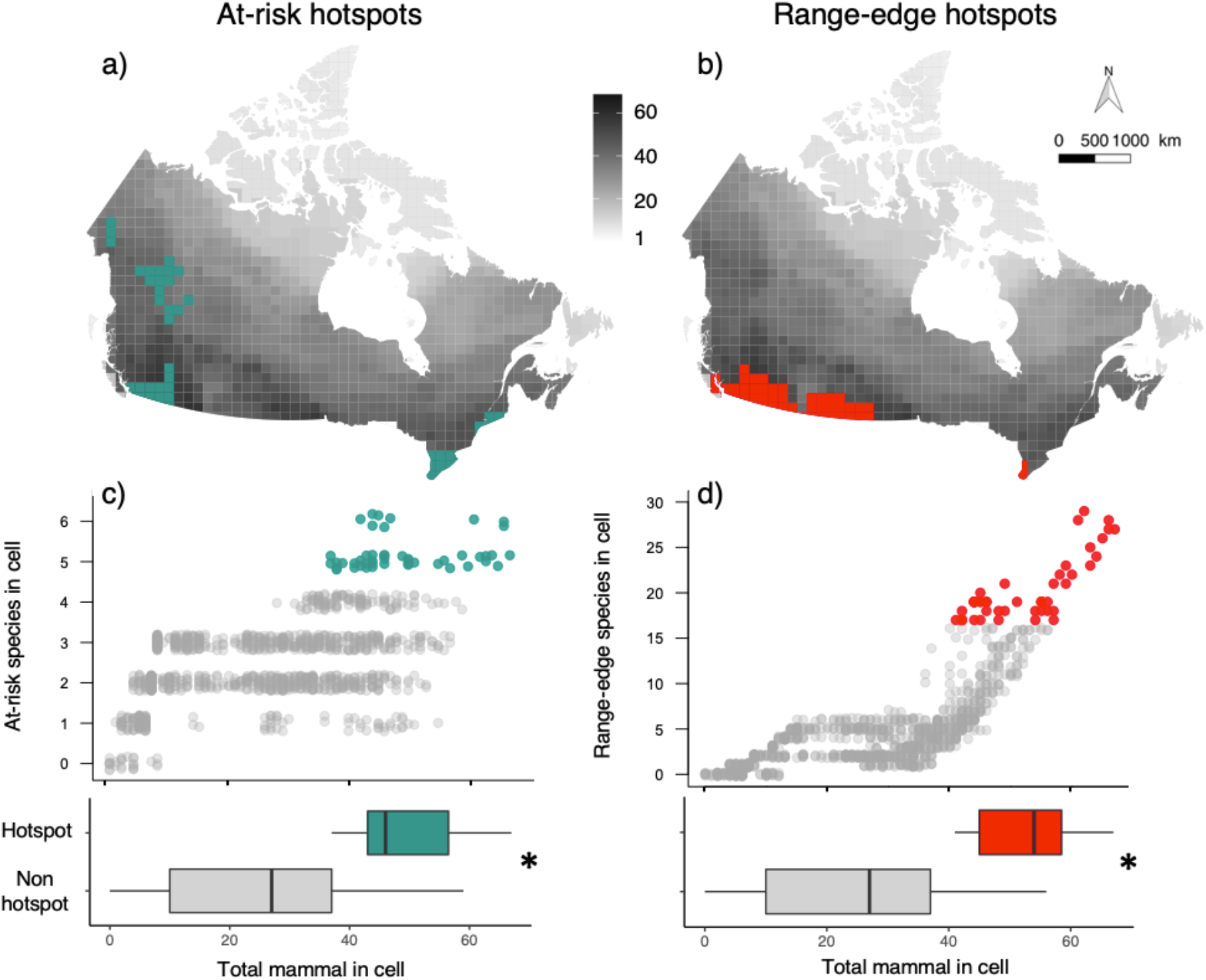
High spatial overlap between hotspots and total mammal richness. Maps of Canada are divided into 1284 100×100 km grid cells, shaded in greyscale to show total species richness of terrestrial mammals. Coloured cells show locations of hotspot cells, i.e. those with the highest richness of a) mammals deemed at-risk in Canada (teal), and b) mammals that only occur in Canada in the northernmost 20% or less of their global distribution (red). (c & d) Relative diversity of hotspot (coloured) vs. non-hotspot (grey) cells. Middle plots show raw data, where each point represents one grid cell. Bottom: Hotspots have higher total mammal richness than non-hotspot cells (* indicates a significant difference within panel). Boxes summarize raw data, showing the median and quartiles, formatting as per Fig. 1.

Whereas regression models assume data points are independent, we detected significant spatial autocorrelation; i.e. hotspot cells for at-risk taxa occurred closer to each other than expected by chance (Moran’s *I*±SD = 0.0628 ± 0.00137, *P* < 0.001), as did range-edge hotspots (Moran’s *I*±SD = 0.116 ± 0.00137, *P* < 0.001). We accounted for this spatial autocorrelation by incorporating the spatial configuration of data via a fixed effect covariate (*s*), using the Spatial Eigenvector Mapping method (Griffith & Peres-Neto 2006; Dormann et al. 2007; Thayn & Simanis 2013). We selected the neighborhood size of the spatial configuration covariate using the value that maximized model fit based on its Akaike Information Criterion (AIC) value (Supporting Information, Augustin et al. 1996; Dormann et al. 2007). Including *s* as a fixed effect corrected the autocorrelation in residuals and improved model fit (Supporting Information).

We tested whether total mammal richness was higher in at-risk hotspots than other cells using a GLM with binomial error structure. Each cell contributed one data point. The cells’ at-risk hotspot status (at-risk hotspot or not) was a binomial response, and total mammal richness and the spatial covariate *s* were fixed effects (at-risk hotspot ~ total richness + *s*). Significance of predictors was determined using likelihood ratio Chisquare tests as for Question 1. This approach is consistent with previous studies comparing at-risk vs. total richness (Prendergast et al. 1993; Orme et al. 2005; Ceballos & Ehrlich 2006), but we recognize two complicating issues: first, that the response and predictors are not fully independent as an increase in at-risk richness automatically increases total richness; second that smaller cells (bisected by the Canadian border) may contain fewer species due to sampling error. We ran two alternate GLMs to test whether our results were sensitive to either issue, one with richness of secure mammals as the predictor (at-risk hotspot ~ secure mammal richness + *s*; Supporting Information), and one including cell area as a fixed effect (Supporting Information). Both models reached the same conclusions as our original model, so we present analyses using total richness and not including cell size to facilitate comparison with previous studies.

Similarly, we tested whether total mammal richness was higher in hotspots of range-edge taxa richness than other cells (range-edge hotspot ~ total richness + *s*). Results using richness of secure mammals instead of total richness were again similar (Supporting Information).

## Results

In Canada, mammal richness is highest toward the south and along the western mountains (Fig. 1a, Fig 2. grey scale maps), although this pattern was not apparent for the 22 taxa deemed at-risk (Fig. 1b). Half (50%) of all ‘Canadian’ mammals had less than 20% of their global range in Canada.

### Q1) How do range area in Canada and range percentage in Canada relate to national conservation status?

Range area in Canada differed significantly between nationally at-risk and nationally secure mammals, but peripherality did not (Fig. 1). As predicted, mammals at-risk of extinction in Canada had smaller Canadian ranges than mammals considered secure (model a: *X^2^*_df=1_ = 6.28*, P* = 0.012; Fig. 1c). Contrary to our predictions, at-risk mammals did not have smaller fractions of their global ranges in Canada (model b: *X^2^*_df=1_ = 0.39, *P* = 0.53; Fig. 1d). Probability of being at-risk in Canada increased as Canadian range area decreased (area: *X^2^*_df=1_ = 4.54*, P* = 0.033), but was not affected by range percentage in Canada after controlling for range area (percentage: *X^2^*_df=1_ = 0.06*, P* = 0.85, model c).

### Q2) Do hotspots of at-risk or range-edge taxa coincide with high overall diversity?

Of the 1285 100×100 km grid cells in Canada, cells with the highest richness of at-risk and range-edge mammals (‘hotspots’) coincided with areas of high total richness (Fig. 2). Hotspots of at-risk mammals had significantly higher total mammal richness than other cells (*X^2^*_df=1_ = 140.3*, P* < 0.0001; Fig. 2c). At-risk hotspot cells each contained 37 to 67 mammal species (24–44% of Canada’s mammal richness; Fig. 2c), and together were home to 80% of the 153 Canadian mammals in our data. Similarly, hotspots of range-edge species, i.e. species only present in Canada in ≤20% of their range, also had significantly higher mammal richness than other cells (*X^2^*_df=1_ = 266.8*, P* < 0.0001; Fig. 2d). Range-edge hotspot cells each contained 41 to 67 mammal species (27–44% of Canada’s mammal richness), and together were home to 81% of the 153 Canadian mammals in our data. The two types of hotspot partially coincided with each other: almost a third of at-risk hotspot cells were also range-edge hotspots and vice versa (15 of 79 hotspot cells overlapped; Fig. 2).

## Discussion

Our quantitative analyses of how mammals are spatially distributed in Canada revealed three main findings. First, our results confirmed earlier qualitative assessments (Yakimowski & Eckert 2007; Gibson et al. 2009) that Canada is a land of edge populations; half the mammals that live in Canada only do so at the northernmost 20% or less of their global range. Second, by quantifying range area we showed that species with smaller Canadian ranges are more likely to be nationally at-risk (Fig. 1a), but that species with smaller fractions of their range in Canada were not (Fig. 1d and model c). Thus we found no evidence that range-edge populations are inherently more vulnerable due to their demography (e.g. putatively small or isolated populations) or coincidence with areas of high human population density. Third, we found significant spatial overlap between high mammal diversity and hotspots of both at-risk mammals and range-edge mammals (Fig. 2). This result is particularly exciting, as it suggests that protecting habitat for at-risk mammals could have significant co-benefits to mammals considered secure, and that protecting range-edge populations need not involve a trade-off with protecting Canada’s overall mammal diversity.

Our finding of high spatial overlap between total mammal richness and at-risk mammal richness (Fig. 2c) contrasts results from other areas and taxa. Globally, hotspots of at-risk mammals do not strongly overlap areas of high total mammal richness (Ceballos & Ehrlich 2006), nor do hotspots of at-risk birds strongly overlap with hotspots of total bird richness (Orme et al. 2005). Diversity is driven largely by energy availability and biogeographic history (Gaston 2000), whereas threats to wildlife are driven by human impacts (Schipper et al. 2008; Szabo et al. 2012). Poor global congruence between at-risk and total diversity suggests these drivers have different global distributions (Orme et al. 2005). In Canada however, mammal richness concentrates in the south and west (Fig. 1), as does the intensity of human land conversion (Kerr & Cihlar 2004; Kerr & Deguise 2004). To the extent that diversity and human impacts co-occur in other high-latitude countries, high overlap between at-risk and total diversity may be more common at regional than global scales.

Of course, habitat protection generally happens on a much smaller scale than our 100×100 km grid cells, and species rarely occupy all land area within their occurrence polygon at fine scales. Protecting habitat within a hotspot may not capture all the species whose ranges overlap with the larger grid cell. Nevertheless, identifying hotspots at a coarse scale is still useful for conservation planning; the hotspots identified here should now be priorities for finer scale assessments (Rodrigues et al. 2004). Further, we are in an era of species on the move, as species shift their geographic distributions in response to anthropogenic change (Chen et al. 2011; Freeman et al. 2018). Protecting habitat in high diversity areas provides habitat options that nearby species may well use in the future, even if they don’t use them currently.

Cells that were hotspots for both at-risk and range-edge taxa are particularly interesting for long term conservation. Since the highest densities of mammals deemed at-risk in Canada occur in the United States (Fig. 1b), one effective conservation strategy for many at-risk taxa in Canada may be to maintain habitat connections to US populations. This would both enable natural immigration and population replenishment, and provide bridges if or when species begin to shift northward in response to warming, both of which could bolster Canadian populations. Indeed, given worldwide range shifts to higher latitudes (Chen 2011), Canada’s current range-edge species may become much less peripheral in the coming decades, and much of Canada’s future biodiversity may be on our doorstep. Protecting habitat in the ‘double hotspots’ is a win-win-win, protecting high numbers of mammals overall, protecting the at-risk species to whom Canada deems it has a national responsibility, and protecting Canada’s future biodiversity in terms of currently peripheral species and the wildlife refugees from climate change coming our way.

## Acknowledgements

Funds for this work were provided by NSERC (Discovery grant to ALH) and the E. Gordon Edward fund in support of undergraduate research in biology at McGill University. We thank Pascale Caissy for her help with QGIS.

## Supporting Information

SI is available online. Appendix S1 = distributions of species assessed by COSEWIC grouped by COSEWIC threat status. Appendix S2 = grid map of Canada. Appendix S3 = results from analyses exploring sensitivity of hotspot models and from alternate analytical decisions described in Methods. Appendix S4 = AICs of spatial GLMs according to different neighborhood distances. The authors are solely responsible for the content and functionality of these materials. Queries (other than absence of the material) should be directed to the corresponding author.

## Appendix S1

**Appendix S1:**
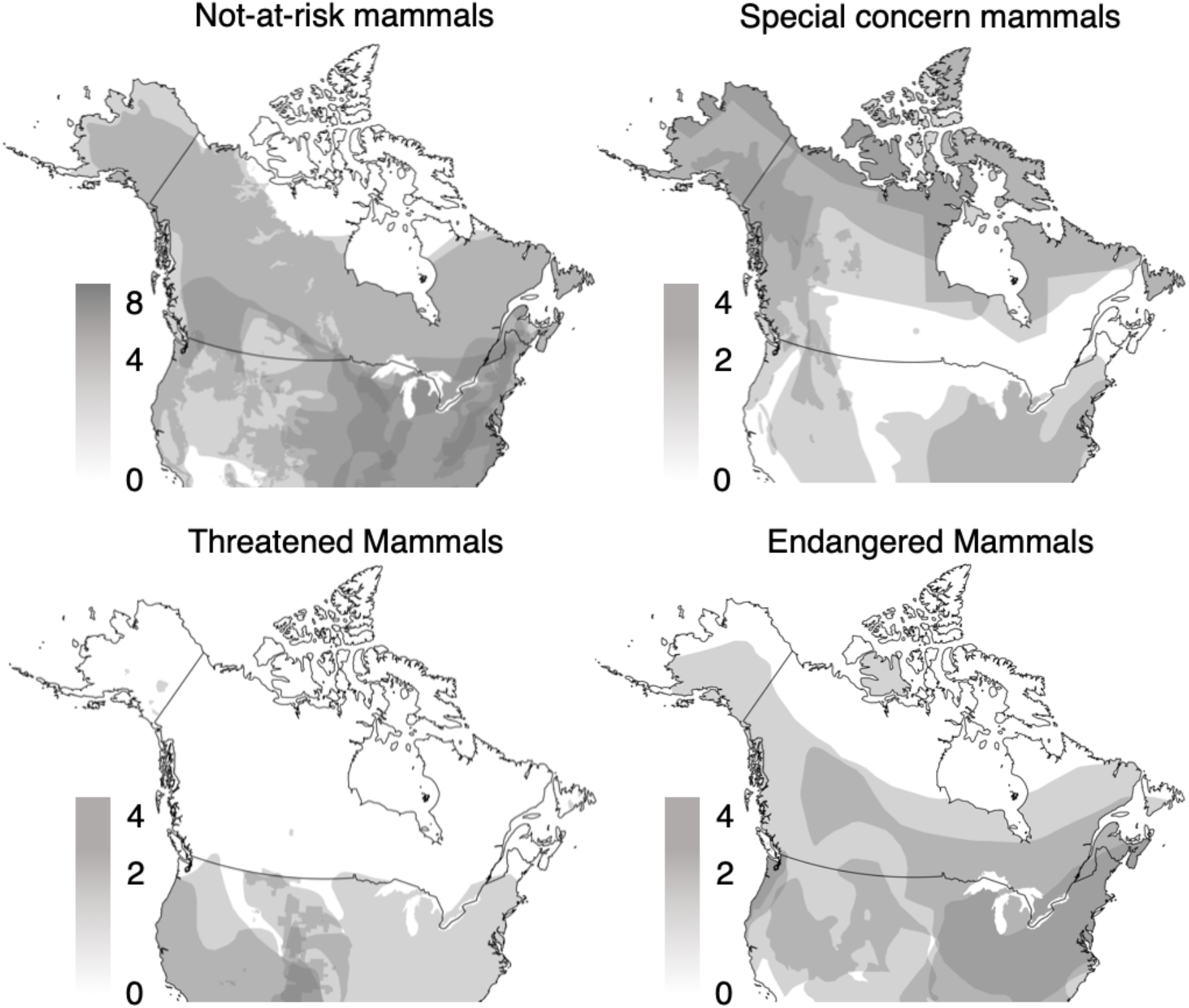
Spatial distribution of the Canadian terrestrial mammals that have been assessed by COSEWIC, separated by their COSEWIC status: Not-at-risk of extinction (n=8), Special concern (n=10), Threatened (n=6), and Endangered (n=8).

## Appendix S2

**Appendix A2:**
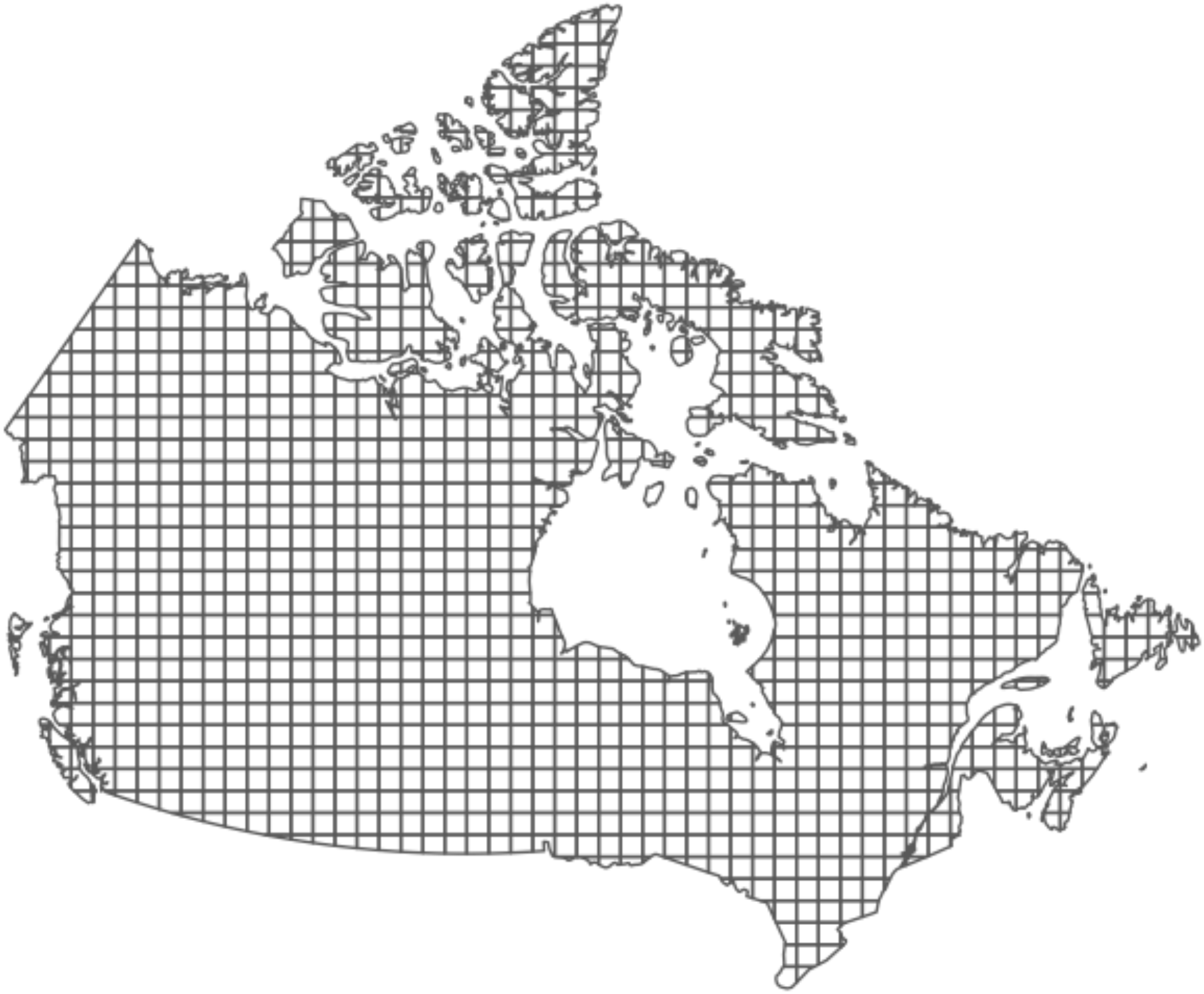
Grid map of Canada. The grid is a 100 x 100 km cell grid overlapped on the map of Canada projected in an Albers equal area projection. Mammal diversity metrics were calculated for each grid cell for analyses in *Question 2* (hotspots).

## Appendix A3

Results from additional models exploring the sensitivity of hotspot analyses. All models compared a binary response (a cell was or was not a hotspot) to a measure of overall richness (fixed effect). Results show the analysis of deviance of each model. Models presented in the main text are in bold. *Spatial covariate*: Including a term to capture the spatial covariance in the data (*s*: Spatial Eigenvector Mapping) improved model fit compared to equivalent models without *s* (models 1 vs 2, models 5 vs 6). Including *s* also corrected the autocorrelation in residuals for the at-risk hotspot model (model 2 without *s*: Moran’s *I* ±SD = 0.046 ± 0.0014, *P* < 0.001; model 1 with *s*: Moran’s *I* ±SD = - 0.00060 ±0.0014, *P* = 0.90) and the range-edge hotspot model (model 7 without *s*: Moran’s *I*±SD = 0.063±0.0014, *P* < 0.001; model 6 with *s*: Moran’s *I*±SD = −0.0023 ±0.0013, *P* = 0.020). *Richness of secure mammals vs. all mammals*: Comparing richness of secure mammals between hotspots vs. non-hotspots yielded similar results to comparing total mammal richness (at-risk + secure) between hotspots vs. non-hotspots for analyses of at-risk hotspots (model 1 vs 3) and range-edge hotspots (models 5 vs 7). *Cell area*: Accounting for cell area in models did not change conclusions (models 1 vs 4, 5 vs 8). *Threshold for being considered a range-edge taxon*: Reducing the threshold for ‘range-edge’ taxa from ≤20% of their global range in Canada to ≤10% of their range in Canada did not qualitatively change results (model 5 vs 9).

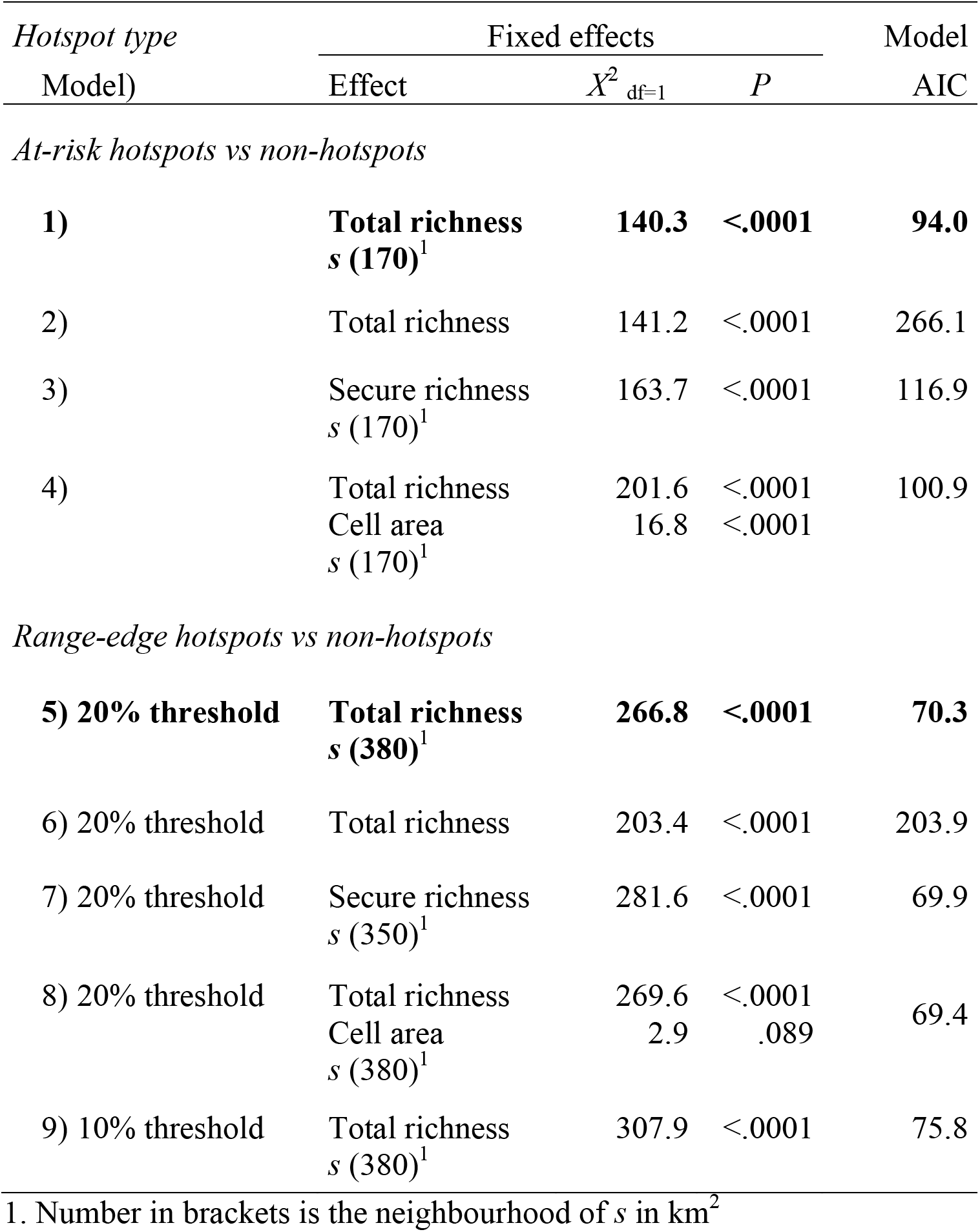

## Appendix S4

**Appendix S4:**
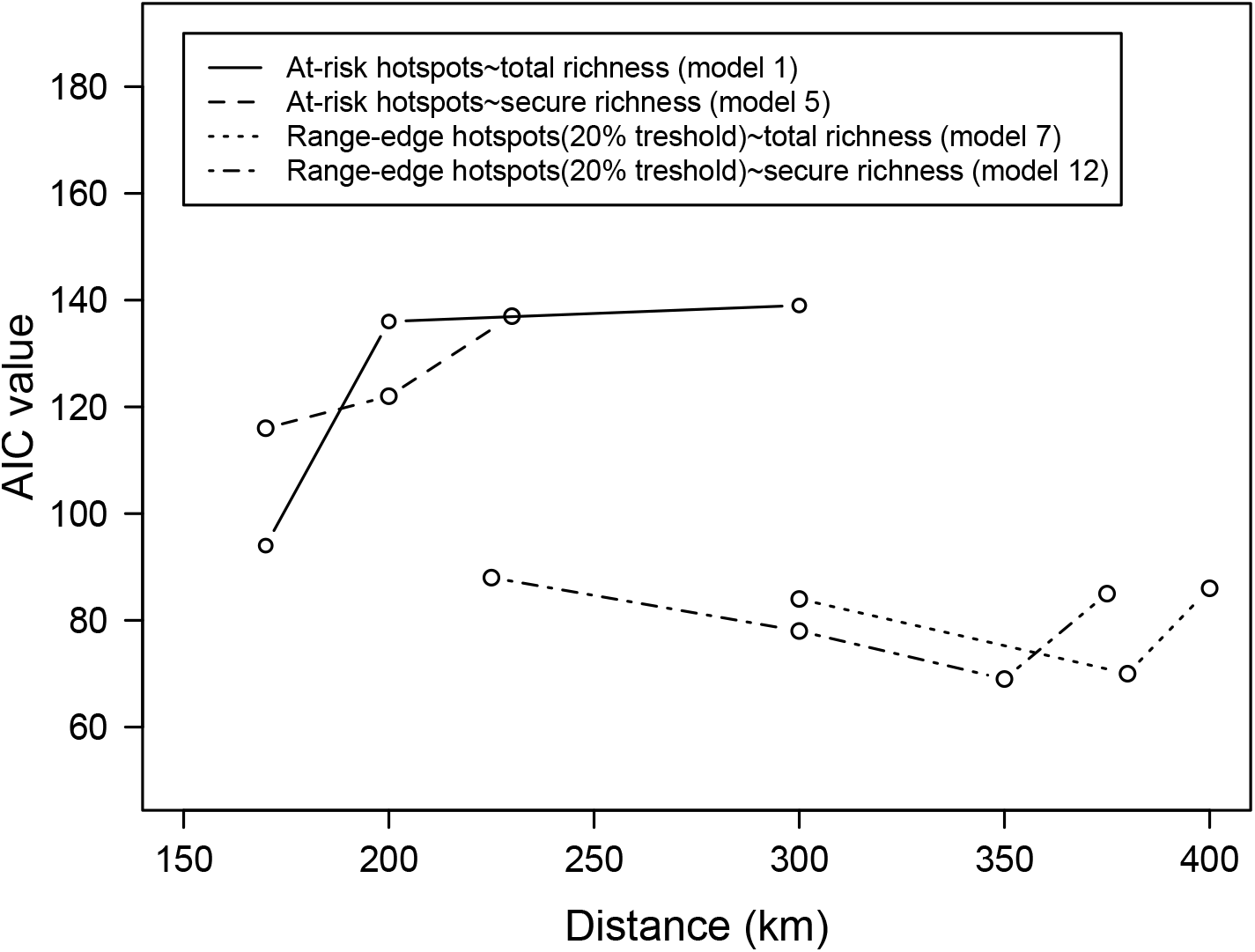
Akaike information criterion (AIC) values for different distances of neighborhood within the spatial extra covariate integrated to GLMs. Neighborhood distances were used to obtain the spatial covariate using Spatial Eigenvector Mapping (SEVM). Absent values indicate distances for which the eigenvector filtering could not yield results. AICs for different distances are presented for all three GLM models presented in the methods. The distance resulting in the lowest AIC was selected for the given model.

## Literature Cited

ArtDatabanken. 2015. The 2015 red list of Swedish species. Uppsala, Sweden.

Augustin N, Mugglestone M, Buckland S. 1996. An autologistic model for the spatial distribution of wildlife. The Journal of Applied Ecology 33:339–347.

Brown JH, Stevens GC, Kaufman DM. 1996. The geographic range: size, shape, boundaries, and internal structure. Ecology 27:597–623.

Buckley LB et al. 2010. Phylogeny, niche conservatism and the latitudinal diversity gradient in mammals. Proceedings of the Royal Society B: Biological Sciences 277:2131–2138.

Ceballos G, Ehrlich PR. 2006. Global mammal distributions, biodiversity hotspots, and conservation. Proceedings of the National Academy of Sciences of the United States of America 103:19374–19379.

Cheffings CM, Farrell L, Dines TD, Jones RA, Leach SJ, McKean DR, Pearman DA, Preston CD, Rumsey FJ, Taylor I. 2005. Species status No. 7. The vascular plant red data list for Great Britain. Peterborough, UK.

Chen IC, Hill JK, Ohlemüller R, Roy DB, Thomas CD. 2011. Rapid range shifts of species associated with high levels of climate warming. Science 333:1024–1026.

Coristine LE, Kerr JT. 2011. Habitat loss, climate change, and emerging conservation challenges in Canada. Canadian Journal of Zoology 89:435–451.

Dormann C et al. 2007. Methods to account for spatial autocorrelation in the analysis of species distributional data: a review. Ecography 30:609–628.

Eckert CG, Samis KE, Lougheed SC. 2008. Genetic variation across species’ geographical ranges: The central-marginal hypothesis and beyond. Molecular Ecology 17:1170–1188.

Freeman BG, Lee-Yaw JA, Sunday JM, Hargreaves AL. 2018. Expanding, shifting and shrinking: The impact of global warming on species’ elevational distributions. Global Ecology and Biogeography 27:1268–1276.

Gaston KJ. 2000. Global patterns in biodiversity. Nature 405:220–227.

Gibson S, Van der Marel R, Starzomski B. 2009. Climate change and conservation of leading-edge peripheral populations. Conservation Biology 23:1369–1373.

Griffith D, Peres-Neto P. 2006. Spatial modeling in ecology: the flexibility of eigenfunction spatial analyses. Ecological Society of America 87:2603–2613.

Hargreaves AL, Eckert CG. 2019. Local adaptation primes cold-edge populations for range expansion but not warming-induced range shifts. Ecology Letters 22:78–88.

Hunter ML, Hutchinson A. 1994. The virtues and shortcomings of parochialism: Conserving species that are locally rare, but globally common. Conservation Biology 8:1163–1165.

IUCN (International Union for the Conservation of Nature). 2018. The IUCN red list of threatened species. Version 2018-1. Gland, Switzerland. Available from http://www.iucnredlist.org (accessed October 11, 2018).

Jenkins CN, Pimm SL, Joppa LN. 2013. Global patterns of terrestrial vertebrate diversity and conservation. Proceedings of the National Academy of Sciences of the United States of America 110:E2603–E2610.

Kerr JT, Cihlar J. 2004. Patterns and causes of species endangerment in Canada. Ecological Applications 14:743–753.

Kerr JT, Deguise I. 2004. Habitat loss and the limits to endangered species recovery. Ecology Letters 7:1163–1169.

Klemet-N’Guessan S, Jackiw R, Eckert CG, Hargreaves A. 2019. Edgy conservation: Canadian at-risk plants are overwhelmingly range-edge populations and under-studied. bioRxiv:682823.

Lawton JH. 1993. Range, population abundance and conservation. Trends in Ecology and Evolution 8:409–413.

Le Saout S et al. 2013. Protected areas and effective biodiversity conservation. Science 342:803–805.

Lesica P, Allendorf FW. 1995. When are peripheral populations valuable for conservation? Conservation Biology 9:753–760.

Mimura M, Aitken SN. 2010. Local adaptation at the range peripheries of Sitka spruce. Journal of Evolutionary Biology 23:249–258.

Myers N. 1988. Threatened biotas: “Hot spots” in tropical forests. The Environmentalist 8:187–208.

Myers N, Mittermeier RA, Mittermeier CG, Fonseca GAB, Kent J. 2000. Biodiversity hotspots for conservation priorities. Nature 403:853–858.

Natural Earth. 2018. Natural earth vector countries. Version 4.1.0. Available from https://www.naturalearthdata.com (accessed September 10, 2018).

Orme CDL et al. 2005. Global hotspots of species richness are not congruent with endemism or threat. Nature 436:1016–1019.

Pebesma E. 2018. Simple features for R: standardized support for spatial vector data. R Journal 10:439–446.

Pironon S, Papuga G, Villellas J, Angert AL, García MB, Thompson JD. 2017. Geographic variation in genetic and demographic performance: new insights from an old biogeographical paradigm. Biological Reviews 92:1877–1909.

Prendergast JR, Quinn RM, Lawton JH, Eversham BC, Gibbons DW. 1993. Rare species, the coincidence of diversity hotspots and conservation strategies. Nature 365:335–337.

R Core Team. 2018. R: A language and environment for statistical computing. R Foundation for Statistical Computing, Vienna, Austria. Available from https://www.r-project.org/.

Rassi P, Hyvärinen E, Juslen A, Mannerkoski I. 2010. The 2010 red list of finnish species. Available from: www.nationalredlist.org (accessed June 2019).

Reid W V. 1998. Biodiversity hotspots. Trends in Ecology & Evolution 13:275–280.

Ricketts TH et al. 2005. Pinpointing and preventing imminent extinctions. Proceedings of the National Academy of Sciences 102:18497–18501.

Rodrigues ASL et al. 2004. Global gap analysis: priority regions for expanding the global protected-area network. BioScience 54:1092.

Sagarin RD, Gaines SD. 2002. The “abundant centre” distribution: to what extent is it a biogeographical rule? Ecology Letters 5:137–147.

Samis KE, Eckert CG. 2009. Ecological correlates of fitness across the northern geographic range limit of a pacific coast dune plant. Ecology 90:3051–3061.

Schipper J et al. 2008. The status of the world’s land. Science 322:225–230.

Schloss CA, Nunez TA, Lawler JJ. 2012. Dispersal will limit ability of mammals to track climate change in the Western Hemisphere. Proceedings of the National Academy of Sciences 109:8606–8611.

Szabo JK, Khwaja N, Garnett ST, Butchart SHM. 2012. Global patterns and drivers of avian extinctions at the species and subspecies level. PLoS ONE 7:1–9.

Thayn JB, Simanis JM. 2013. Accounting for spatial autocorrelation in linear regression models using spatial filtering with eigenvectors. Annals of the Association of American Geographers.

Thomas C et al. 2004. Extinction risk from climate change. Nature 427:145–148.

Van Rossum F, Vekemans X, Gratia E, Meerts P. 2003. A comparative study of allozyme variation of peripheral and central populations of Silene nutans L. (Caryophyllaceae) from Western Europe: Implications for conservation. Plant Systematics and Evolution 242:49–61.

Venables W, Ripley B. 2002. Modern applied statistics with S. Fourth ed. Springer, New York. Available from http://www.stats.ox.ac.uk/pub/MASS4.

Yakimowski SB, Eckert CG. 2007. Threatened peripheral populations in context: Geographical variation in population frequency and size and sexual reproduction in a clonal woody shrub. Conservation Biology 21:811–822.

